# Identifying Putative Pathogenic Non-Coding Variants in Unresolved Rare Disease Patients Using Topologically Associated Domains

**DOI:** 10.64898/2026.09.17.752339

**Authors:** Anthony M. Gacita, Matthew Pahl, Manuel Diz Torres, Shiva Ganesan, Justin J. Blair, Khushbu Patel, Rajagopalan Ramakrishnan, Laura Conlin, Ingo Helbig, Struan F.A. Grant

## Abstract

Unresolved rare disease is a major public health challenge affecting ∼300 million people worldwide. At least 50% of these individuals remain genetically unresolved after applying exome sequencing and/or whole genome sequencing. One source of these missing diagnoses is the presence of rare variants within the non-coding genome that are detected but not interpreted by whole genome sequencing. In order to systematically evaluate candidate pathogenic non-coding variants, we created the <u>G</u>enomic <u>A</u>nalysis of <u>V</u>ariants in <u>U</u>nresolved <u>R</u>are <u>D</u>isease (GAVURD) system. GAVURD leverages trio whole genome sequencing alignment data to produce a short list of putative pathogenic non-coding variants for a given proband. GAVURD uses best practices for *de novo* and rare inherited variant identification, links variants to human disease genes harnessing topologically associated domain (TAD) data, and rank prioritizes variants based on phenotypic overlap. As a proof-of-concept, we applied GAVURD to ten probands with unresolved rare disease and implicated six potentially causal non-coding variants based on a confluence of evidence supportive of pathogenicity. The GAVURD system serves an important role in prioritizing candidate non-coding causal variants for unresolved rare disease that can serve as the high value and informed focus of additional functional follow-up studies.

## INTRODUCTION

Rare diseases refer to diseases that affect fewer than 1 in ∼2,000 people. While individually rare, in total, rare diseases affect ∼300 million people worldwide and represent a major public health challenge. Genetic factors are widely thought to contribute to at least ∼70% of rare disease.[1, 2] However, at least 50% of rare disease patients that undergo genetic testing have no genetic cause defined for their symptoms (i.e. are unresolved).[3, 4] A molecular diagnosis is vital to the clinical care of rare disease patients as it can provide disease-specific prognostication, and identify target variants for the increasing number of DNA-modifying therapies currently under development. Therefore, increasing the diagnostic rate in rare disease will positively affect the health of millions of individuals worldwide.

The sources of missing diagnoses in unresolved rare disease probands are numerous. Poorly understood environmental effects and complex polygenic inheritance patterns likely contribute, but pinpointing the exact etiologies is challenging. In contrast, a major underexplored source of missing diagnoses is the non-coding genome. Clinical whole genome sequencing (WGS), which can identify variants throughout the entire genome, is analytically targeted to the coding portion the human genome. Most clinical variant analysis pipelines are currently unable to interpret non-coding variants. However, there are established examples of pathogenic non-coding variants that can cause Mendelian disease including those causing polydactyly by affecting enhancer for *SHH* or isolated pancreatic agenesis by affecting an enhancer for *PTF1A*.[5–7] In addition, a case of aniridia was shown to be driven by a single point mutation in an enhancer located ∼150kb upstream of *PAX6*, the master regulator of eye development.[8] Similarly, a variant in an enhancer of *TBX5,* a key heart development transcription factor, was identified as a cause of congenital heart disease.[9] Likewise, variants disrupting a tissue-specific regulatory element in *HK1* cause congenital hyperinsulinism. [10] However, these were discovered via low throughput intensive studies at individual loci. Therefore, improving the systematic interpretation of non-coding variants for Mendelian disease represents an opportunity to improve the diagnostic rate in rare disease.

The non-coding genome has multiple functions including regulating gene expression, gene splicing, transcript stability, and the various functions of non-coding RNAs.[11] A major challenge in non-coding variant interpretation is the high volume of such variants in a typical proband. However, in rare disease diagnostics, the amount of potential pathogenic variants can be narrowed by focusing on *de novo* variants, i.e. those that arose newly in a proband. Calling *de novo* variants from trio WGS data is challenging due to interference from sequencing errors. Many bioinformatic tools have been developed to identify *de novo* variants from trio WGS data including machine learning and consensus-based approaches.[12–14] These tools have various levels of accuracy and require validation, which is a requirement for a translational approach.

Once candidate pathogenic variants are identified in the non-coding genome, predicting their functional impact is a challenge. Epigenetic and functional genomic data can provide clues on regulatory variant function, but alone are often insufficient for clinical non-coding variant interpretation. Another approach is to compare the clinical phenotype of the proband with the phenotype of patients who have mutations in the coding regions of genes nearby a given non-coding variant. This approach was popularized by Exomiser, which uses a semantic similarity between gene and proband Human Phenotype Ontology (HPO) terms to rank coding variants.[15] The same team extended this analysis to the non-coding genome with Genomiser.[16] Using simulated genomes, the authors report that Genomiser can successfully recover a known pathogenic non-coding variants in 77% of probands. These results indicate the power of this approach and have been useful for projects aiming to improve the diagnostic rate in rare disease such as the Undiagnosed Disease Network.[17] An important limitation is the reliance on simulated genomes, which insert a known pathogenic non-coding variant into a background of benign variants. The generalizability of this approach to identify novel pathogenic variants in actual trio whole genome sequencing data remains an unanswered question.

In this paper, we sought to develop and evaluate a tool that combines the power of variant calling with phenotype-based gene prioritization. The ‘genomic analysis of variants in unresolved rare disease’ (GAVURD) system represents an end-to-end tool that combines variant calling with gene-based ranking. To achieve this goal, we implemented a consensus based *de novo* variant and rare inherited homozygous variant calling algorithm. The identified variants can then be analyzed to implicate known human disease genes that reside with the same topologically associated domain (TAD) as the given variant. Finally, GAVURD ranks putative pathogenic variants by the strength of the phenotypic overlap between the non-coding variant’s implicated gene and the patient’s phenotype using Human Phenotype Ontology (HPO) terms. As such, GAVURD operates on WGS alignment data, a ped file describing sample relationships, and HPO term lists. We applied the optimized GAVURD system to 10 genetically unresolved rare disease probands and identified a limited, high value set of candidate pathogenic non-coding variants that can serve as targets for additional functional validation. The GAVURD system represents a useful tool that serves the need to improve the diagnostic yield of WGS in rare disease patients.

## RESULTS

### Informatic Pipeline to Identify Candidate Pathogenic Variants from WGS Data

To identify candidate pathogenic variants in genetically unresolved rare disease probands, we first developed a bioinformatic pipeline to call *de novo* and rare homozygous inherited variants from trio WGS data. We focused on *de novo* and rare homozygous inherited variants given that many known highly penetrant pathogenic non-coding variants are *de novo* or homozygous in probands and very rare in the population. This strategy allows for the filtering of variants based on inheritance and population information, without the need for functional genomic data. The pipeline starts with alignment files from trio WGS data and calls variants with three pipelines-GATK, DeepTrio, and DRAGEN. (**Fig. 1**) The resulting variant calls are compared, and only variants called by at least 2 of the 3 callers are kept as consensus variants. Variants present in structurally complex regions (Encode blacklists, segmental duplications, low mappability regions) are removed. Only variants that are rare in the general population and local population are retained as the final variant set.

**Figure 1.**
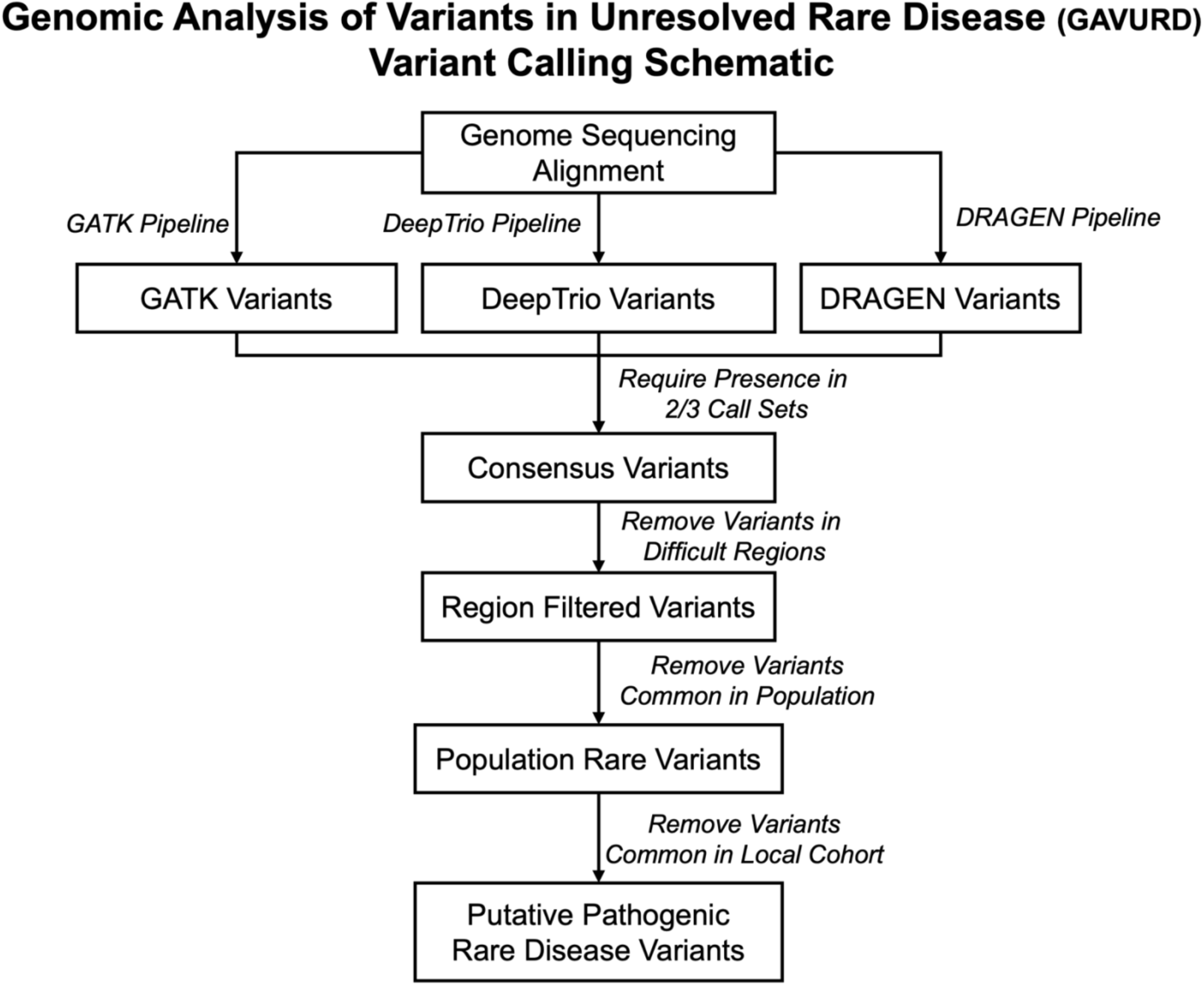
Schematic of GAVURD variant calling pipeline. Genome sequencing alignment files are analyzed with three pipelines-GATK, DeepTrio, and DRAGEN. Only variants called by 2/3 pipelines are retained. Final variant set is created by removing variants in difficult regions and variants common in the general or local population.

Identifying *de novo* variants from trio WGS data is known to be challenging, as only 40-90 true *de novo* variants are expected per proband.[18] However, given the hundreds of variants initially produced by *de novo* variant callers, the majority of new variants initially detected in a proband are likely the result of sequencing errors. Multiple approaches have been developed to maximize *de novo* variant calling accuracy. One previous approach that utilizes consensus across multiple variant callers performed well [14]; as such, we developed our own consensus approach that combines GATK, DeepTrio, and DRAGEN variant callers. To determine appropriate variant quality control (QC) filtering metrics, we evaluated our pipeline with ‘genome in a bottle’ (GIAB) data, which includes high confidence *de novo* variant calls.[19] We graphed the distribution of multiple variant QC metrics including genotype quality, sequencing depth, alternative allele depth, and allele balance for true *de novo* and false positive *de novo* variants. (**Supplemental Fig. 2**) We determined QC cutoffs that prioritized the recovery of true positives at the cost of increased false positives for each variant caller. (**Supplemental Table 1)** Using these cutoffs, our pipeline achieved a recall of 0.98 and precision of 0.93 for GIAB *de novo* variants. (**Supplemental Fig. 1**)

As part of our variant calling pipeline, we also sought to identify homozygous inherited variants (subsequently referred to as inherited variants). We defined an inherited variant as a variant that is heterozygous in parents and homozygous in the proband. Calling inherited variants has higher confidence due to independent evidence of variant calls in multiple samples. For consistency, we modified the *de novo* variant QC filtering metrics to appropriate values for inherited variants. Probands have many more inherited variants than *de novo* variants, so to limit the number of inherited variants found by our pipeline, we excluded variants that were common in the general and/or local population, as these are unlikely to explain rare genetic disease.

### WGS Analysis from 10 Unresolved Rare Disease Probands Identifies *De Novo* and Inherited Variants

We leveraged trio WGS from 10 randomly selected unresolved rare disease probands who were sequenced at the Children’s Hospital of Philadelphia (CHOP). We selected probands who had undergone clinical exome or WGS that was non-diagnostic. The phenotypes of these probands included individuals with VACTERL (vertebral, anorectal, cardiac, tracheal, esophageal, renal, and limb defects) sequence, congenital diaphragmatic hernia, Hirschsprung’s disease and multiple patients with complex congenital heart disease, consistent with the largely unknown genetic etiology associated with these congenital disorders.[20–22] (**Supplemental Table 4)**

The GAVURD variant calling pipeline was used to identify *de novo* variants in these 10 genetically unresolved probands. Results indicated that individual variant callers using GIAB-optimized quality filters separately produced a high number of false-positive *de novo* variants per proband-219.4 single nucleotide polymorphism (SNPs), 167.2 SNPs, and 183.1 SNPs for GATK, DeepTrio and DRAGEN respectively. (**Fig. 2**) Additional analysis including a consensus comparison step, removing variants in difficult regions and allele frequency filters yielded in an average of 72.6 *de novo* SNPs and 8.3 *de novo* indels per proband. Analysis of these variants indicated that the chromosomal location of variants largely correlated with chromosomal length, as expected for randomly distributed variants. (**Supplemental Fig. 3A**) *De novo* variants also showed significant enrichment (p-value < 0.001) for GC dinucleotide sequences. (**Fig. 4B**) This result is consistent with the known increased rate of CG to TG *de novo* mutations due to the spontaneous deamination of CpG nucleotides.[23] The distribution of *de novo* variants per proband shows a range of 53 – 109 variants, which is consistent with the known human *de novo* mutation rate.[24] (**Fig. 4A**) Overall, these findings indicate that the consensus-based GAVURD *de novo* variant caller identifies true *de novo* mutational events.

**Figure 2.**
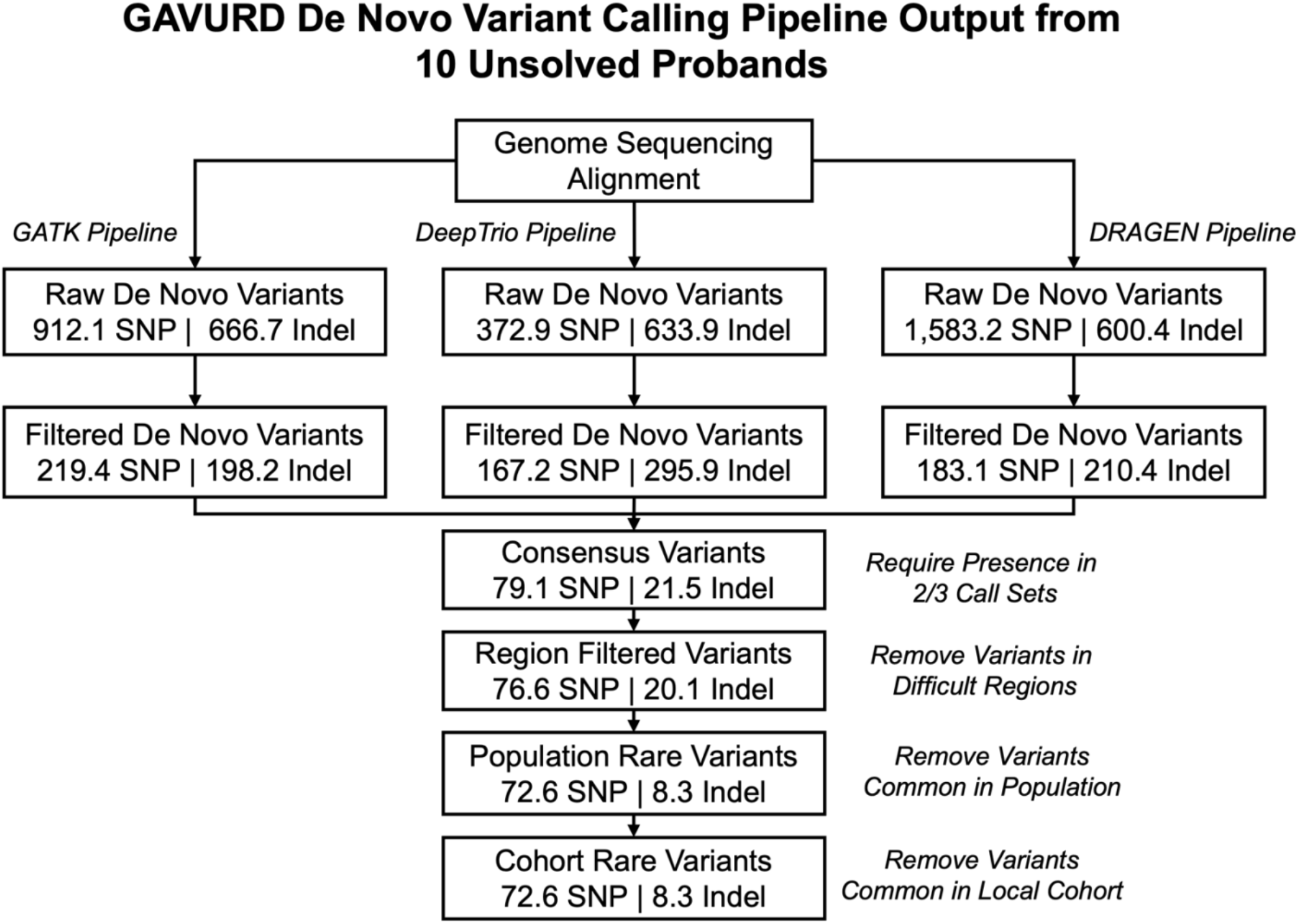
GAVURD de novo variant calling pipeline output from 10 unsolved rare disease probands. Average number of de Novo SNPs and indels are indicated at each step. Quality control (QC) and consensus comparison steps provide the strongest filters. The GAVURD de novo variant calling pipeline identifies an average of ∼72 de Novo SNPs and ∼8 de Novo indels per proband.

The GAVURD variant calling pipeline also evaluated 10 unresolved probands for inherited variants. As expected, the number of inherited variants per proband is much larger than *de novo* variants. After a consensus and region filtering step, 168,396.7 inherited SNPs and 22,070 inherited indels per proband were observed. (**Fig. 3**) However, removing variants common in the general population or local population, significantly reduced the number of inherited variants to 92.4 SNPs and 8.6 indels per proband. The power of this filter is likely two-fold: rare variants are unlikely to be heterozygous in unrelated parents and sequencing errors (especially indels around repeat sequences) are likely overrepresented in population/local sequencing databases. In contrast to *de novo* variants, inherited variant chromosomal location are not consistent with random locations as there is not a clear association with chromosome length. (**Supplemental Fig. 3B**) The distribution of inherited variants per proband also indicates a broader distribution ranging from 46 to 347 inherited variants/proband. (**Figure 4A**) It is possible that probands with higher numbers of inherited variants show higher rates of identity by descent due to parental shared ancestry. Another contributing factor is that variants common in probands with ancestries not well represented in population databases are more likely to survive filtering due to a low allele frequency in the databases. Indeed, the proband with 347 inherited variants reported Hispanic ancestry. Overall, these findings demonstrate the power of using population and local allele frequency to prioritize inherited variants that are more likely to be causative of rare disease.

**Figure 3.**
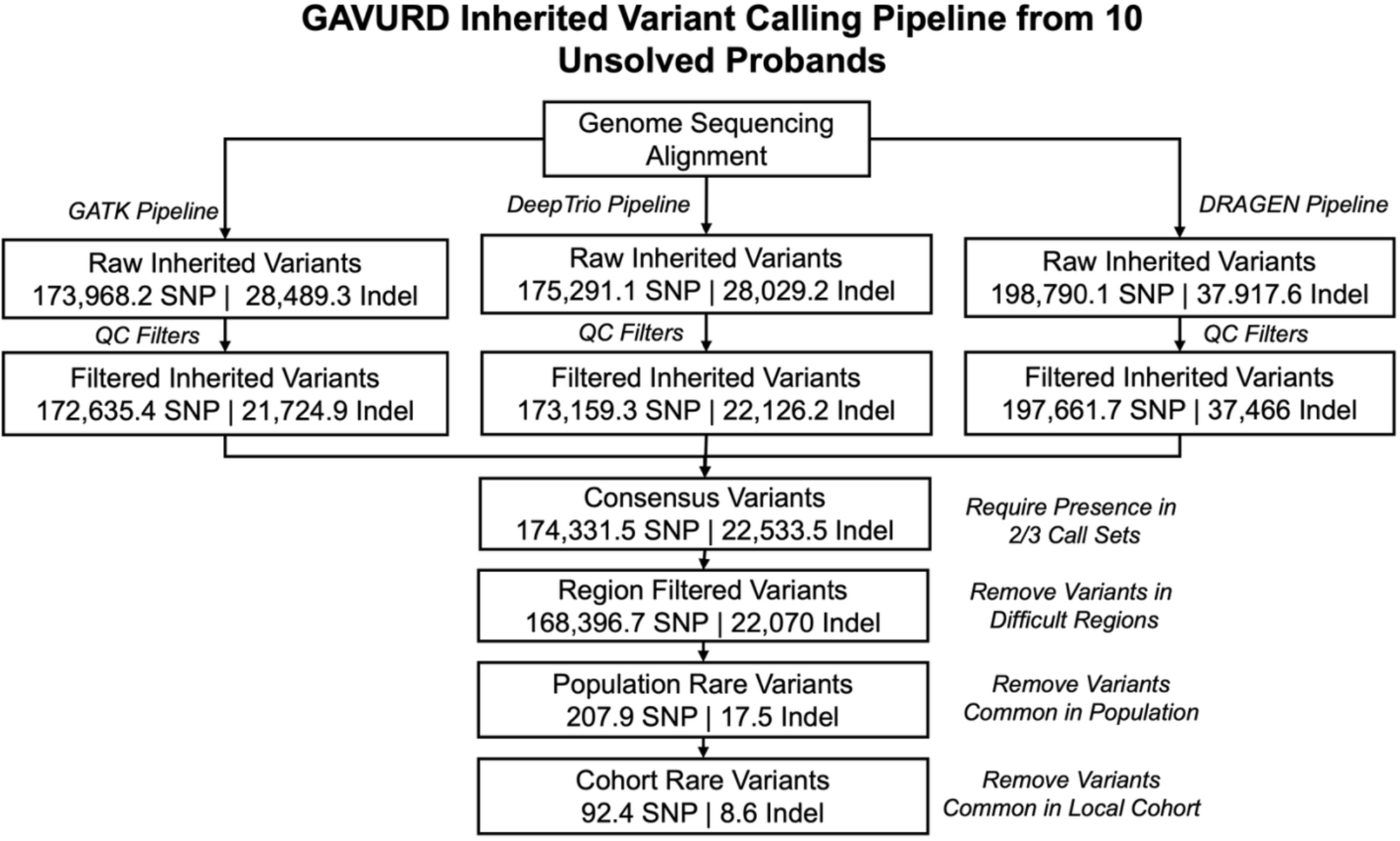
GAVURD inherited variant calling pipeline output from 10 unsolved rare disease probands. Average number of inherited SNPs and indels are indicated at each step. Population and local cohort allele frequency metrics provide the strongest filtering. The GAVURD inherited variant calling pipeline identifies an average of ∼92 inherited SNPs and ∼9 inherited indels per proband.

### Implicating Human Disease Genes Using Topologically Associated Domain and Phenotype Data

In total, the GAVURD variant analysis identified a total of 181.6 variants per proband that are either *de novo* or rare inherited variants. Variant annotation showed that variants were preferentially located in intronic, intergenic, and non-coding RNA sequences. **(Fig. 4C & 4D)** This is consistent with a random placement of variants as intronic and intergenic sequences make up the largest fraction of the human genome. We manually reviewed all high effect coding variants (stop gain, missense, and splicing) and did not identify any variants expected to cause disease, which confirmed these patients’ previously non-diagnostic clinical testing.

**Figure 4.**
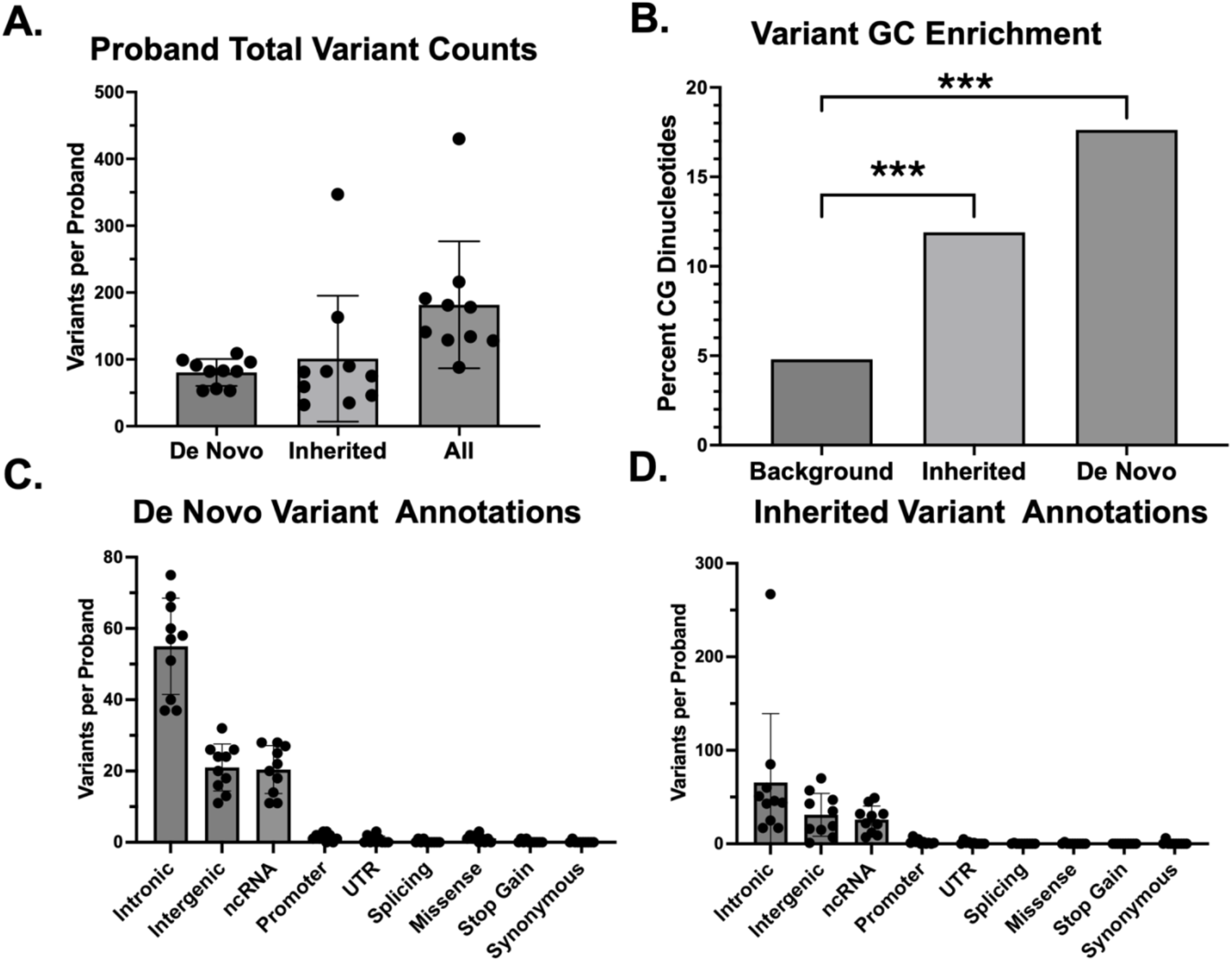
Features of GAVURD called variants from 10 unsolved rare disease probands. **A.** Distribution of the number of de novo and inherited variants called by the GAVURD variant calling pipeline. **B**. Assessment of CG Dinucleotide enrichment demonstrates that de novo and inherited variants are significantly more likely to occur at CG sites. **C.** Variant annotations relative to MANE transcripts for de Novo variants shows intronic, intergenic, and ncRNA predominant annotations. **D.** Variant annotations relative to MANE transcripts for inherited variants shows intronic, intergenic, and ncRNA predominant annotations. *** p < 0.001, binomial test

Given the lack of clear disease-causing variants and enrichment for intergenic/intronic regions, we developed a system to prioritize variants based on their likelihood to regulate to known human disease genes. The non-coding genome harbors important regulatory sequences that can act over long distances to affect gene expression. Regulatory sequences interact with gene promoters through three-dimensional folding that is organized into topologically associated domains (TADs). TADs represent the fundamental structure of genome organization, and non-coding variants are more likely to regulate a gene within a shared TAD i.e. the search space for the affected gene should be within the boundaries of a given TAD. Indeed, previous work has indicated that TAD data can aid in interpretation of non-coding copy number variants.[25]

We gathered in house and publicly available Hi-C data from 17 different tissue and cell lines representing cell types across the body. We identified TAD boundary locations supported by the Hi-C interaction data. Comparison of TAD boundaries across samples demonstrated that similar tissues tended to have boundary locations that cluster together. (**Supplemental Fig. 4A**) The distribution of TAD boundaries supported by multiple samples indicated a large population of boundaries found in a single sample and revealed a population that was shared across many samples. (**Supplemental Fig. 4B**). Analysis of CTCF ChIP-seq data from ENCODE showed that boundaries supported by multiple samples had higher levels of CTCF binding. **(Supplemental Fig. 4C**) These results provide higher confidence in the multi-sample TAD boundaries as TAD boundaries are generally conserved across tissues. [26] We defined TAD boundaries present in 2 or more samples as more conserved and those present in 9 or more as most conserved boundaries.

Using the most conserved TAD boundaries as a guide, we sought to annotate each variant with a possible target gene. (**Fig. 5A**) The content of the TAD containing a given variant was analyzed for any known human disease genes that share that TAD. Our approach prioritized dominant disease genes for *de novo* variants and recessive disease genes for inherited. When compared to using a linear distance-based boundary (i.e. an arbitrary 1Mb window on each side of the variant site), the most conserved TAD boundary approach implicated significantly fewer genes. (**Fig. 5B**) To determine which genes had the highest likelihood of being causative of a given patient’s phenotype, we integrated phenotypic matching using human phenotype ontology (HPO) terms. We utilized the previously validated Exomiser phenotypic matching system, which has been previously validated.[15] For each implicated gene, we calculated an overlap score between the patient’s HPO terms and the gene’s HPO terms. Results indicated that many genes have little to no phenotypic overlap with the proband, but some probands carry variants that are co-located with genes that could explain their phenotype. (**Fig. 5C**) This process generated a list of putative pathogenic variants ranked by implicated gene’s phenotypic overlap. Top implicated genes, their associated variant, inheritance pattern, and human disease phenotype for the multiple unresolved rare disease probands are listed in **Table 1**.

**Figure 5.**
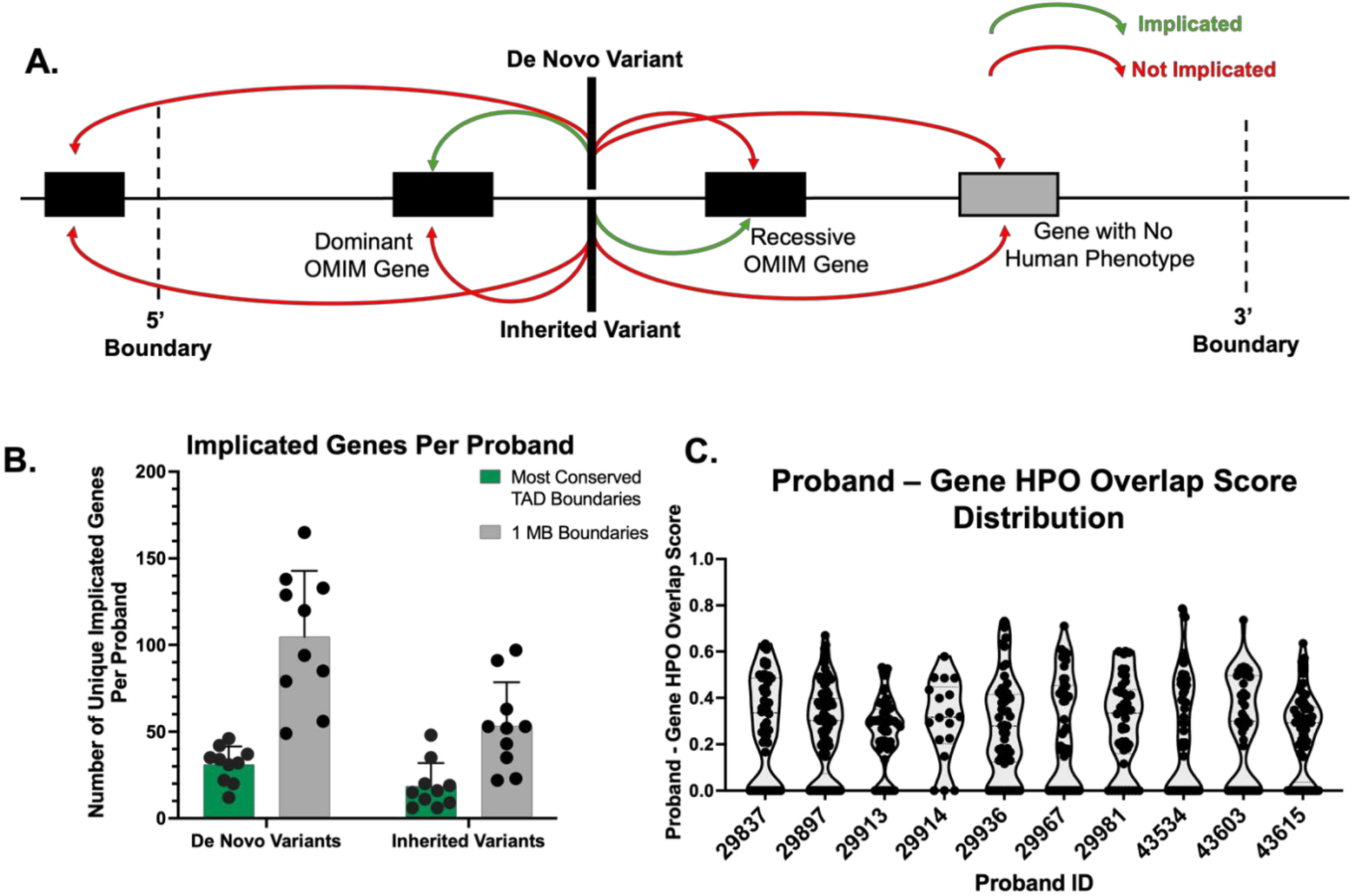
Putative pathogenic non-coding variants implicate genes matching patient phenotypes. **A.** Schematic demonstrating how putative pathogenic non-coding variants implicate disease genes. Variants are mapped to a region based on conserved TAD boundaries. De novo variants implicate dominant human disease genes and inherited variants implicate recessive human disease genes within the same TAD. Only genes with known human disease associations (present in OMIM) are considered. **B.** Comparison of the number of genes implicated when using conserved TAD boundaries or 1MB intervals centered on the variant. Utilizing TAD boundaries decrease the number of genes implicated per proband. **C**. Distribution of the HPO overlap scores between proband HPO terms and gene HPO terms for implicated genes. OMIM, online mendelian inheritance in man. TAD, topologically associated domain. HPO, human phenotype ontology.

**Table 1:**
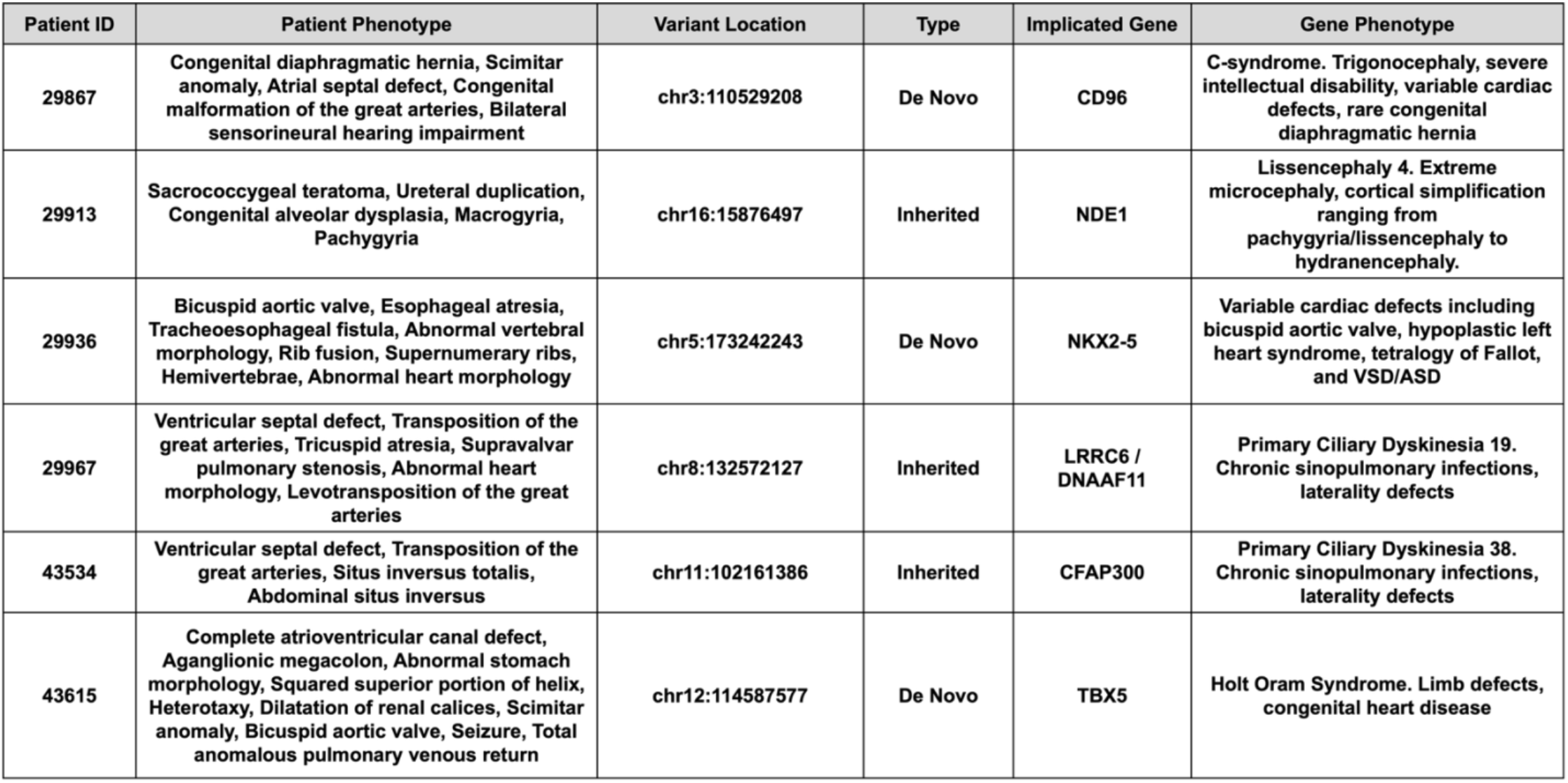
Top scoring putative pathogenic variants in unsolved rare disease probands. Variants were identified by the GAVURD platform and manually evaluated for relevance to patient’s phenotype.

### GAVURD Pipeline Identifies Multiple Putative Pathogenic Non-Coding Variants Implicating Human Disease Genes

The GAVURD variant calling and gene analysis steps generate a ranked list of putative pathogenic variants. We manually reviewed this list to determine the likelihood of pathogenicity. Of the 10 analyzed patients, 6 patients were found to have variants that were causative candidates for their disease. (**Supplemental Table 5**) We discuss two of the strongest candidates below.

Patient 29967 was born with significant congenital heart disease that includes a ventricular septal defect, L-transposition of the great arteries, tricuspid atresia, and pulmonary stenosis. The GAVURD pipeline identified a homozygous inherited variant within the 3’UTR of the *LRRC6/DNAAF11* gene, which is associated with primary ciliary dyskinesia 19. This disorder is associated with chronic sinopulmonary infections as well as laterality defects (abnormal left to right embryonic patterning).[27] The association between congenital heart disease, specifically transposition of the great arteries, and literality defects has been well established. [28] The nominated variant within the 3’UTR creates a miRNA seed sequence that can serve as a binding site for multiple known miRNAs. (**Fig. 6A**) Two of those miRNAs, miR-329-3p and miR-362-3p, are expressed in multiple tissues including the brain, heart, and liver. (**Fig. 6B**) This cumulative data suggests that this rare variant creates a miRNA binding site in the 3’UTR of the LRRC6/DNAAF11 gene, which results in bi-allelic loss of mRNA expression, resulting in ciliary dysfunction, abnormal patterning, and congenital heart disease.

**Figure 6.**
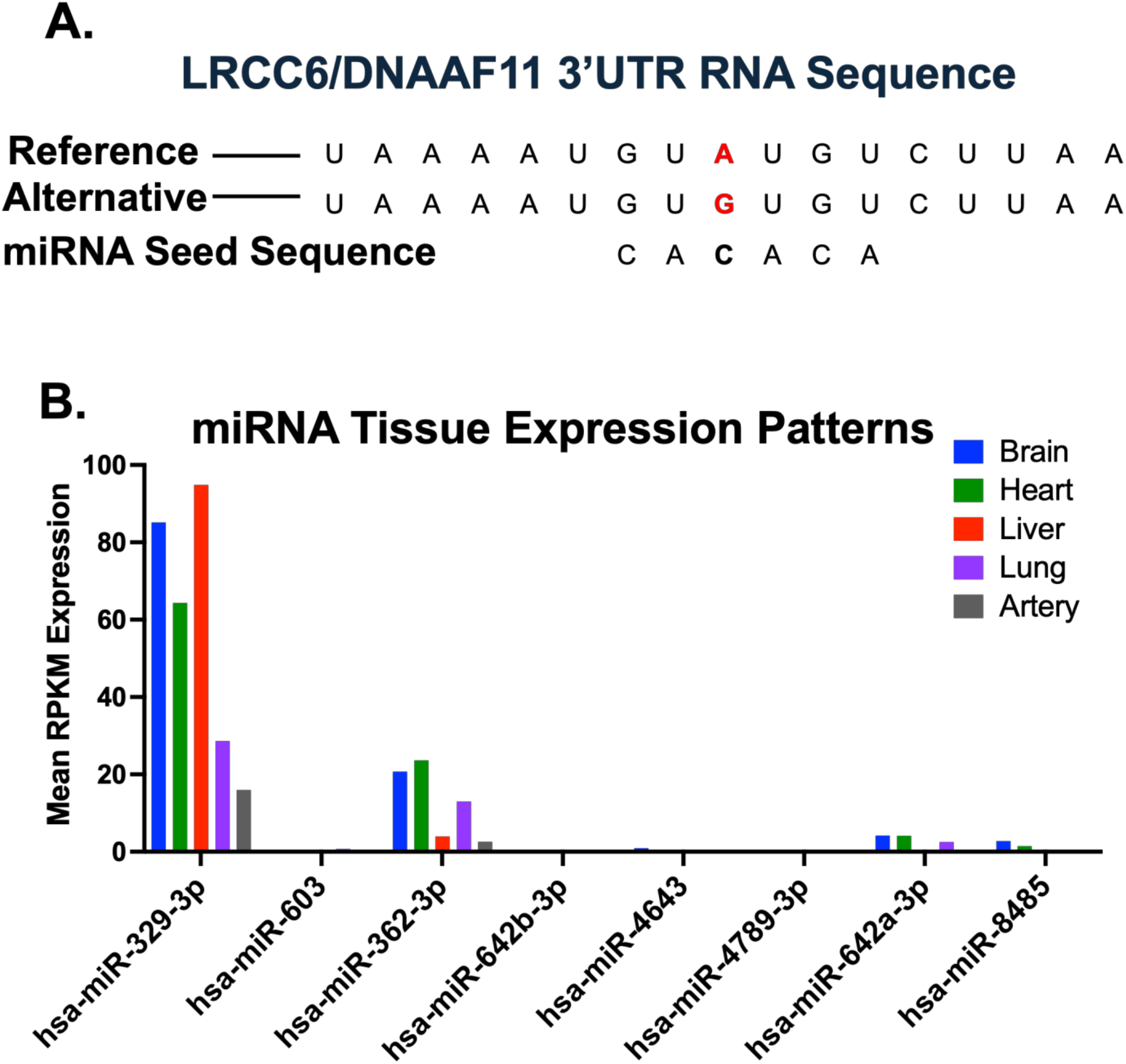
miRNA binding site analysis of the rare inherited variant identified in the 3’UTR of LRCC6/DNAAF11. **A.** The reference and alternative sequences of the LRCC6/DNAAF11 3’UTR are shown. The variant creates a new miRNA seed sequence that is predicted to bind the alternative allele. **B.** Tissue expression of miRNAs with the created seed sequence indicated in A derived from the miRNA tissue expression atlas.

Patient 29936’s presentation includes esophageal atresia, tracheoesophageal fistula, abnormal vertebrae, and bicuspid aortic valve which is most concerning for VACTERL association. The genetic cause of VACTERL association is largely unknown.[20] We observed a *de novo* variant upstream of the gene *NKX2-5*, whose protein product has been associated with multiple types of congenital heart disease, including bicuspid aortic valve.[29] This variant disrupts a highly conserved nucleotide within a *SMAD4* motif. (**Fig. 7B**) *SMAD4* is a master regulator of cardiac development and is known to regulate *NKX2-5.*[30] There is also evidence of *ZBTB20* and *ZNF184* binding at this variant site. (**Fig. 7A)** *ZBTB20* has been shown to protective in cardiac remodeling after myocardial infarction.[31] In contrast, *ZNF184* has no known role in cardiac function or development.

**Figure 7.**
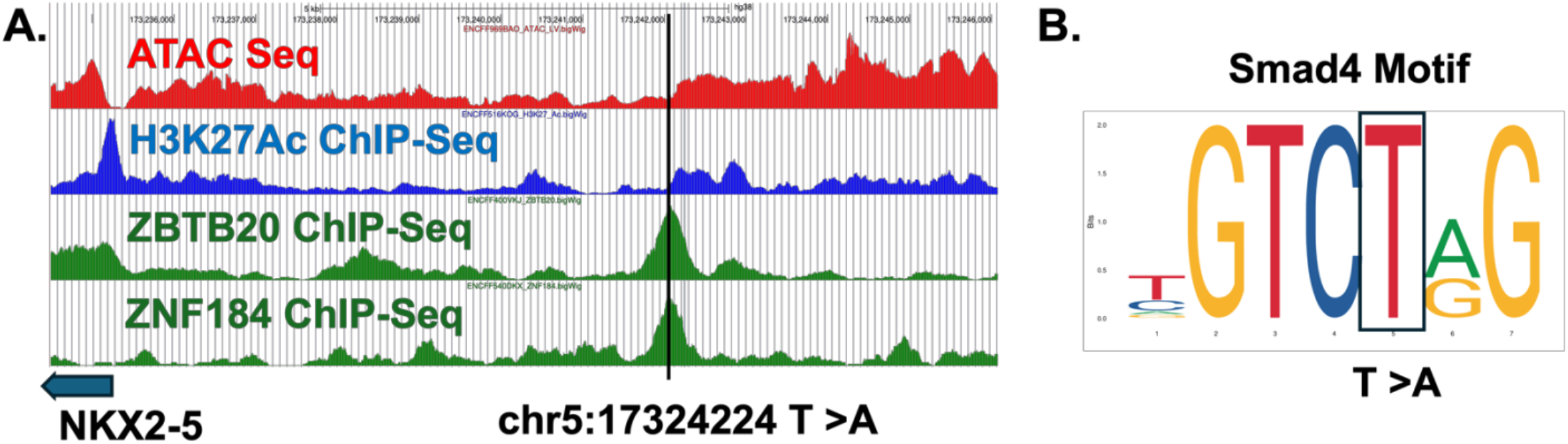
Analysis of epigenetic data overlapping the identified de novo variant upstream of NKX2-5. **A.** Genome browser image showing open chromatin (ATAC-seq), histone modifications (H3K27Ac), and transcription factor binding (ZBTB20 & ZNF164) overlaying the variant site. **B.** Image of the Smad4 binding site demonstrating that the identified variant disrupts a highly conserved site within the binding motif.

In summary, the GAVURD pipeline implicated a non-coding variant with potential disease association in 6 out of 10 cases. The prioritized variants implicate genes that when mutated, result in clinical features overlapping the patient’s phenotype. It is possible these regulatory variants could disrupt gene expression in a tissue specific manner (through transcription factor expression, miRNA expression, etc.), which could cause a tissue-restricted phenotype, but not the entire phenotype.[32] The variants prioritized by GAVURD should be treated as high value candidate non-coding pathogenic variants, where additional evaluation could aid interpretation if these non-coding variants are truly pathogenic.

## METHODS

### Unresolved Rare Disease Proband Cohort Assembly

Rare disease probands were identified through the Birth Defects Biorepository (BDB) Study at the Children’s Hospital of Philadelphia. The BDB study is an IRB-approved protocol (IRB #18-105525) that provides a mechanism to store and provide access to biological specimens and longitudinal clinical, research, and genomic data to support and enhance future research on children with birth defects. All parents provided written informed consent to the use of their and their child’s deidentified data and test results for future research on children with birth defects. The BDB performs research-based trio WGS. To exclude probands with known pathogenic variants, only probands that underwent clinical genome sequencing that resulted as non-diagnostic were included. The BDB study team provided the clinical testing data for participants. For this study, we randomly selected 10 probands as a proof-of-concept study.

### Genome in a Bottle *De Novo* Variant Calling Optimization

To optimize the GAVURD *de novo* variant calling pipeline, we use a truth set *de novo* variants from Genome in a Bottle (GIAB). We downloaded this truth set of *de novo* variants which was derived from the GIAB Ashkenazi Jewish trio, from a previously published study. [13] We included all variants in this set regardless of “manual validation” status. We also removed any truth variants that were not within the callable regions bed files for each trio member, as recommended by GIAB. We removed truth variants that were present in hard-to-call regions and were common in gnomAD (same parameters as final pipeline). This resulted in a truth set of 1,025 *de novo* SNPs and 117 *de novo* indels.

We used the standard GAVURD variant calling pipeline to call *de novo* variants on whole genome sequencing data from the GIAB Ashkenazi trio that was generated by the CHOP clinical genetic testing laboratory. Raw sequencing data was aligned with DRAGEN and the resulting cram files were analyzed with GATK, DeepTrio, and DRAGEN variant calling algorithms with identical settings to the final GAVURD pipeline. We then graphed the distribution of variant quality metrics (genotype quality, read depth, allele balance, alternative allele depth) for true *de novo* (in the downloaded truth set) and false positive *de novo*’s (not in the downloaded truth set). (**Supplemental Fig. 2**) Using these distributions, we empirically determined filtering cutoffs that would keep all true positive variants-maximizing recall. We did not use filter using quality metrics that demonstrated no meaningful separation between true and false positive *de novo* variants. The values for *de novo* variant filtering metrics are indicated in **Supplemental Table 1**.

Previous studies have demonstrated that conducting a force call step where *de novo* variants are force called in parental samples, can identify false positives with read evidence in parents.[14] We conducted a force calling analysis but did not see a significant separation between GIAB true and false positive variants. This result may be due, in part, to the artificial nature of GIAB *de novo* variants. We included the force calling step in our final pipeline but did not use this filter to remove potential false positive *de novo* variants. Instead, variants are tagged with a force calling tag if there is evidence of the variant in parental samples.

We also tested a *de novo* cluster filter that was also supported by previous studies.[14] Clusters of *de novo* variants are a well described phenomenon.[33] However, clusters of variants that extend for long distances are more likely to be technical artifacts. We assessed the GIAB truth set *de novo*’s for variants that were within 10 base pairs of each other and found that the maximum total span of these variants was 11 bp. Therefore, we included a cluster analysis step in the GAVURD pipeline tags clusters of *de novo* variants (within 10 bp of each other) where the total span is > 11bp. Similar to the force calling analysis, the cluster analysis was not used to remove potential *de novo* variants, but tag variants if they are within a long cluster.

### GAVURD Variant Calling Pipeline

The GAVURD variant calling pipeline is designed to identify putative disease variants in trio whole genome sequencing (WGS) alignment data for rare disease probands. We gathered raw sequencing data from 10 rare disease probands. Raw sequencing data was aligned to the hg38 human reference genome using DRAGEN with default settings. Alignment files were used as inputs for GATK, DeepTrio, and DRAGEN variant calling. For GATK variant calling, variants were called with haplotype caller on individual samples and then joint called across each trio. GATK was executed according to standard germline variant analysis including VQSR but without a genotype quality refinement step.[34] DeepTrio was executed according to recommended settings. [33] DRAGEN variant calling was completed in pedigree mode.[35] Variants within the sex chromosomes were called separately by GATK and DRAGEN ensuring appropriate sample and region-specific ploidy arguments (i.e. hemizygous for non-PAR X chromosome variants in male samples).

GATK or a custom script was used identify *de novo*/inherited variants and to apply GIAB optimized variant-caller specific quality control filters to each call set. (**Supplemental Table 1**) The final call sets were compared with bcftools isec and variants called in 2/3 callers were kept. For sex chromosome analysis, variants were required to be called by both GATK and DRAGEN. Variants in hard to call regions were removed with bedtools. To filter variants based on population allele frequency, variants were annotated with gnomad v4 joint allele frequency values. For local allele frequencies, each variant was joint called across the entire 10 sample cohort using GATK. *De novo* variants were kept if gnomAD AF_joint was < 0.01, gnomAD nhomalt_grpmax_joint <= 1, local cohort allele count <= 2, and the local cohort allele number represented at least 90% of callable alleles. Inherited variants were kept if gnomAD nhomalt_grpmax_joint <= 1, the number of homozygous probands in the local cohort was <=2, the number of homozygous non-probands in the local cohort was 0, the number of heterozygous carriers in the local cohort was <= 25, and the local cohort allele number represented at least 90% of callable alleles. For sex chromosome *de novo* variant calls, sex-specific allele frequencies were used in gnomAD.

### GAVURD Variant Caller GC Enrichment Analysis

To determine the enrichment of GC variant calling, we assessed the percentage of *de novo* or inherited variants that occurred at a CpG site. This percentage was compared to the percentage obtained when the variant locations are randomly distributed in the calling intervals with hard to call regions excluded. The difference between inherited and *de novo* variants and background levels was evaluated with a binomial test in R to produce p-values.

### Topologically Associated Domain (TAD) Analysis

Hi-C data generated using the Arima kit or downloaded representing 17 cell types across various tissues. (**Supplemental Table 3**) HiC data was processed using the HiCUP pipeline (v0.7.4) with the restriction enzyme set to Arima kit setting (^GATC/ G^ANTC).[36] Valid reads were subsequently converted to contact 4KB contact matrices using pairtools and cooler.[37, 38]. Cooler merge was used to combine sample replicates. HiTAD was used to call 0-level TAD boundaries for each individual sample using standard settings.[39] We compared the 0-level TAD boundaries across each sample to determine similarity. TAD boundaries were considered identical between samples if they occurred at the same genomic location +/− 12 kb. Sample-wide comparisons were used to generate a multi-dimensional scaling (MDS) plot and distribution of shared boundaries. Boundaries present in at least 2 samples were considered “more conserved” and boundaries present in at least 9 samples were considered “most conserved”. We also generated tissue-specific boundary files.

To evaluate the overlap between TAD boundaries and CTCF binding, we downloaded CTCF peaks from ENCODE ChIP-seq data (encRegTfbsClusteredWithCells_hg38_bed.gz). [40] This data aggregates CTCF ChIP-seq data across approximately 129 cell types. We filtered out CTCF ChIP-seq peaks that overlapped the ECODE blacklist regions (0.77% of peaks).

Overlapping CTCF peaks were merged into non-overlapping intervals for comparison with the boundary files. Each boundary recurrence level (1-17) was compared to the CTCF peak file using bedtools intersect.[41] to obtain the observed percent overlap. We generated a negative distribution for each recurrence level by selecting random genomic intervals that were length matched to the original boundaries and excluded from the Encode blacklist regions 200 times. Enrichment was calculated by comparing the difference between the percent overlap between the original boundary file and the average percent overlap of the negative distributions.

### Implicating Human Disease Genes with GAVURD Gene Analysis

To link variants identified by the GAVURD variant calling pipeline to disease genes, we developed the GAVURD gene analysis pipeline. This process evaluates genes that are within the same TAD as *de novo* or rare inherited variants. This tool uses the “most conserved” TAD boundaries derived from Hi-C data. We focused on known human disease genes using the online inheritance in man (OMIM) database. Implicated genes are tagged based on known inheritance pattern. *De novo* variants implicate genes with dominant inheritance patterns and inherited variants implicate genes with recessive inheritance patterns. Genes with multiple disease associations with different inheritance patterns were kept if any inheritance pattern matches the variant type.

Implicated genes are then prioritized based on Human Phenotype Ontology (HPO) term overlap. The HPO terms associated with each known human gene were downloaded from OMIM. HPO terms for probands were gathered from three sources: manually inputted terms when clinical sequencing was ordered, ctakes natural language processing of clinical notes, and problem list ICD9 to HPO term matching. The three sources of HPO terms were manually evaluated by a clinical geneticist to generate a harmonized list of HPO terms. (**Supplemental Table 4**) We created a custom implementation of the Exomizer HPO scoring tool, hiPHIVE, to evaluate semantic similarity between proband and gene HPO terms.[15] This analysis used the human phenotype only scores to be most conservative. Implicated genes are ranked by hiPHIVE score and the top 10 are outputted for manual analysis.

## DISCUSSION

In this study, we present the genomic analysis of variants in unsolved rare disease (GAVURD) system that implicates putative pathogenic non-coding variants in rare disease probands. This system evaluates WGS data for *de novo* and rare inherited variants and then links those variants to genes with high phenotypic overlap. (**Fig. 8**) We applied this approach to 10 genetically unresolved rare disease cases with congenital anomalies and identified multiple variants with putative pathogenic mechanisms. We note that this tool alone does not generate sufficient evidence for clinical variant classification, but it does produce high-value testable hypotheses to further evaluate with functional studies.

**Figure 8.**
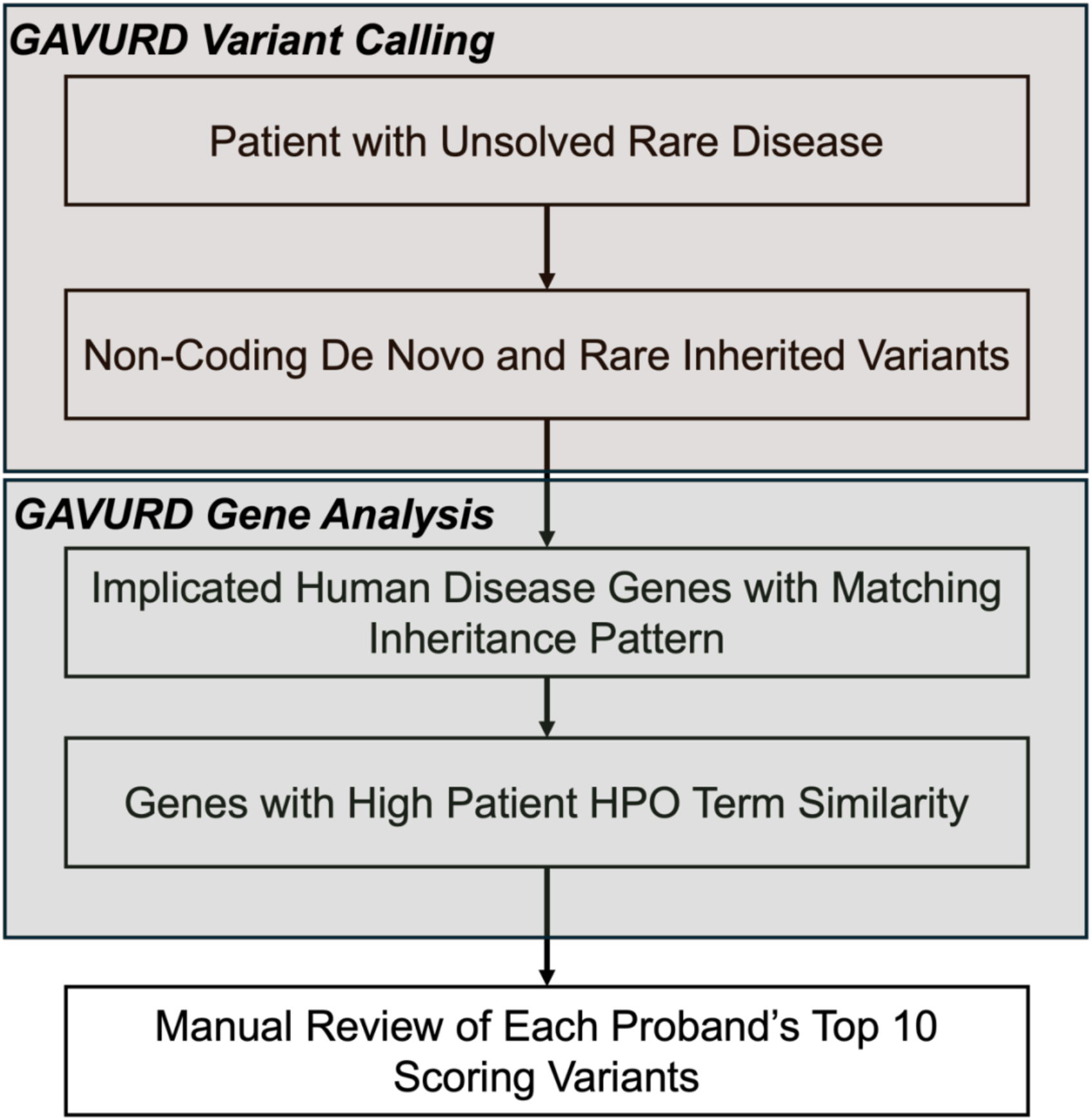
Schematic of the Genomic Analysis of Variants in Unresolved Rare Disease (GAVURD) system. Variant calling highlights variants with genetic evidence for disease association (de novo or rare inherited). Gene analysis prioritizes variants implicating human disease genes with patient phenotypic overlap.

The GAVURD variant calling pipeline utilizes a consensus-based approach to improve the accuracy of *de novo* variant identification. This data adds to the body of work that this is an effective approach for *de novo* variant identification.[14] In addition, we provide variant QC metrics optimized with genome in a bottle variant truth sets. Our approach is simple, automated, and can be scaled to large numbers of unresolved rare disease trios. The GAVURD gene analysis pipeline uses a novel approach that integrates chromatin conformation data to limit a variant’s implicated genes based on TAD coordinates, which is vital given that pathogenic non-coding variants can act over great linear distances. Previously published tools that conduct a similar prioritization strategy, like Genomiser, do not include variant calling in their analysis pipeline.[17] Therefore, GAVURD represents an integrated alignment file-to-prioritized variant list system that can be an important part of the evaluation of undiagnosed rare disease probands.

The GAVURD system has important limitations. The pipeline operates under the assumption that a single high-effect variant is responsible for a proband’s phenotype. While this may be true for many unresolved rare disease probands, it is likely that some unresolved cases are due to complex polygenic and/or environmental mechanisms. The GAVURD pipeline also only considers implicated genes that have been previously linked to human disease. Therefore, it is not suitable for novel gene discovery. Finally, GAVURD inherited variants are required to be homozygous in the proband and ignores the possibility of compound heterozygosity.

## ACKNOWLEDGEMENTS

We would like to thank members of the CHOP Birth Defects Biorepository team: Stacy Woyciechowski, William Gaynor, and Monica Molina. We would also like to thank members of the CHOP ARCUS team, including Gregory Barren for his support. Finally, we thank all the patients and their families for participating in this research study.

This work was supported by an internal CHOP Omics Data Utilization Grant, the Daniel B. Burke Endowed Chair for Diabetes Research, and the Division of Genetics T32 Training Grant.

---

**Supplemental Figure 1.**
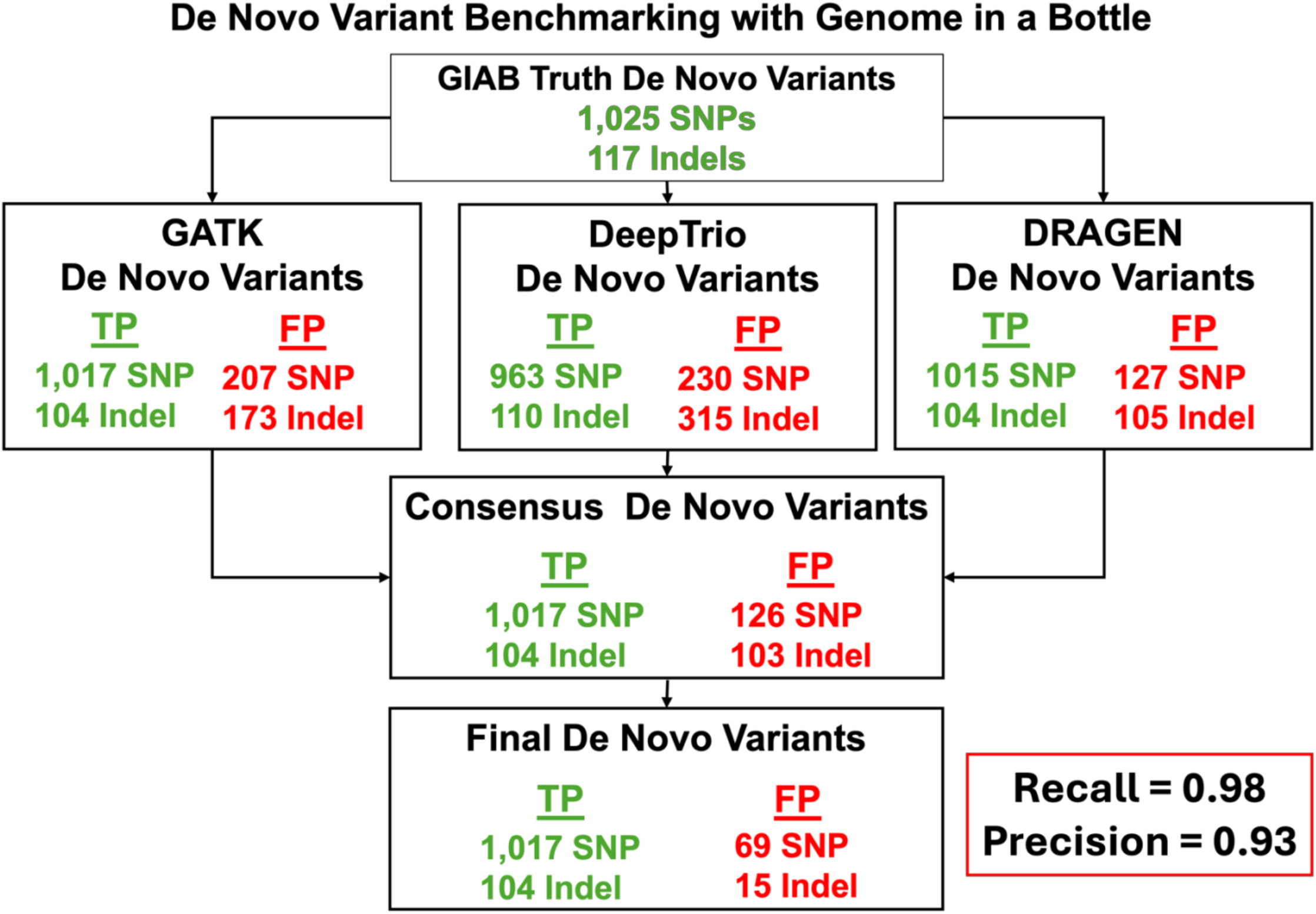
Benchmarking de novo variant calling with Genome in a Bottle (GIAB). Truth set variants were downloaded from GIAB and processed with the GAVURD pipeline. Variant recovery is shown at each step. Final variant counts represent a high recall (0.98) and high precision (0.93).

**Supplemental Figure 2.**
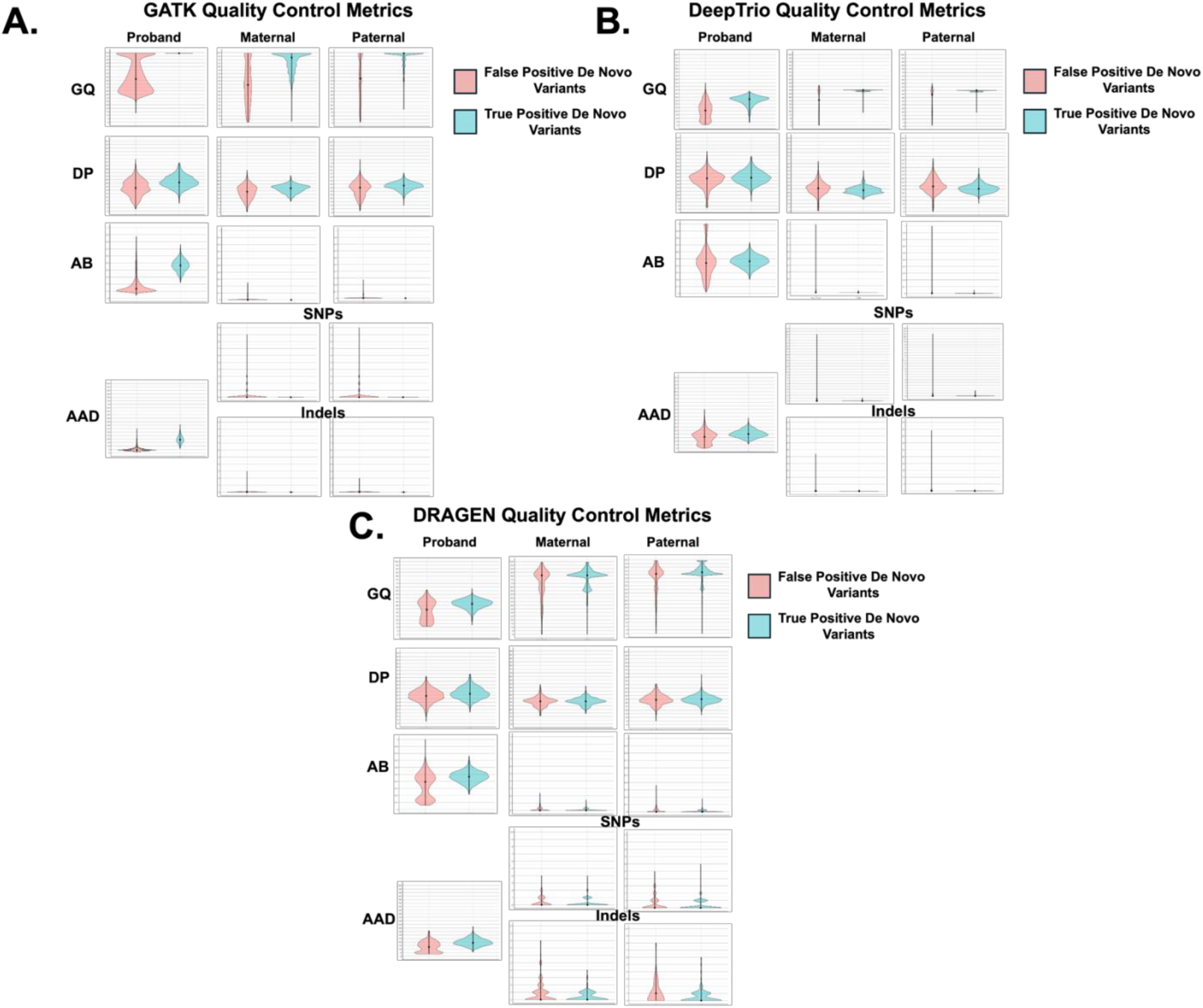
Genome in a Bottle variant metric distributions comparing true positive and false positive de novo variants. **A.** Distribution for GATK called variants. **B.** Distribution for DeepTrio called variants. **C.** Distribution for DRAGEN called variants. GQ, genotype quality. DP, read depth. AB, allele balance. AAD, alternative allele depth.

**Supplemental Figure 3.**
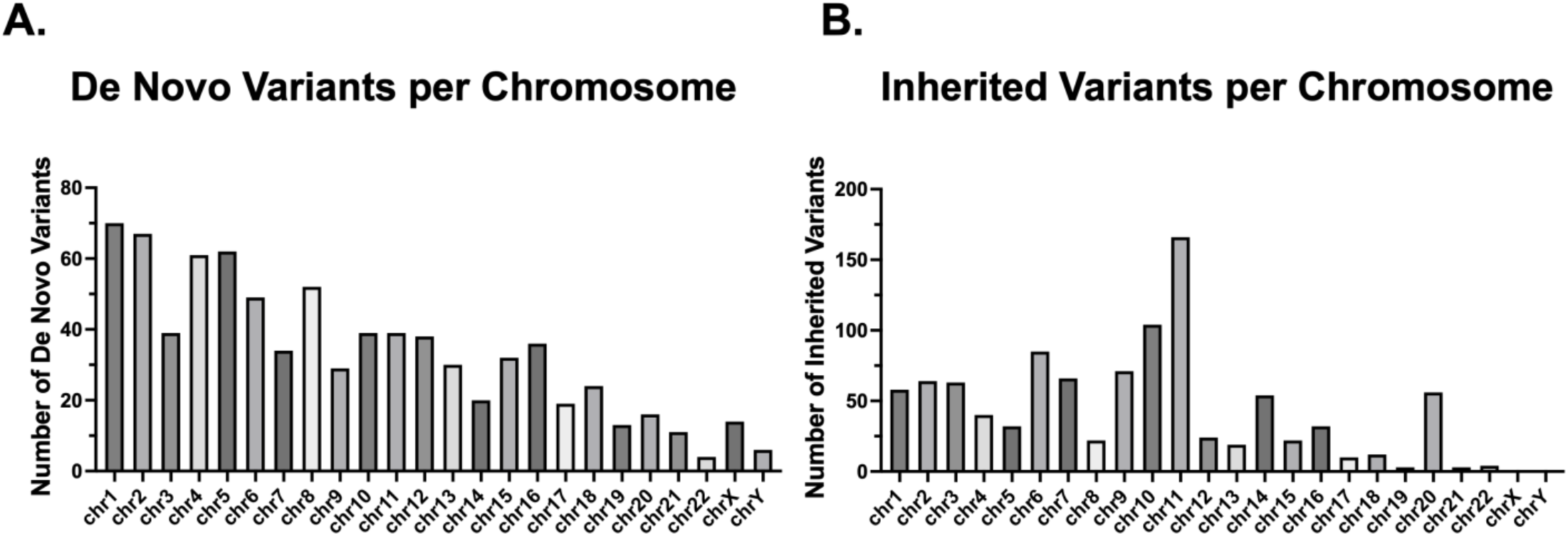
Variants from the 10 unsolved probands presented as counts per chromosome. **A.** De novo variant counts indicating an association with chromosome length. **B.** Inherited variant counts indicating minimal association with chromosome length.

**Supplemental Figure 4.**
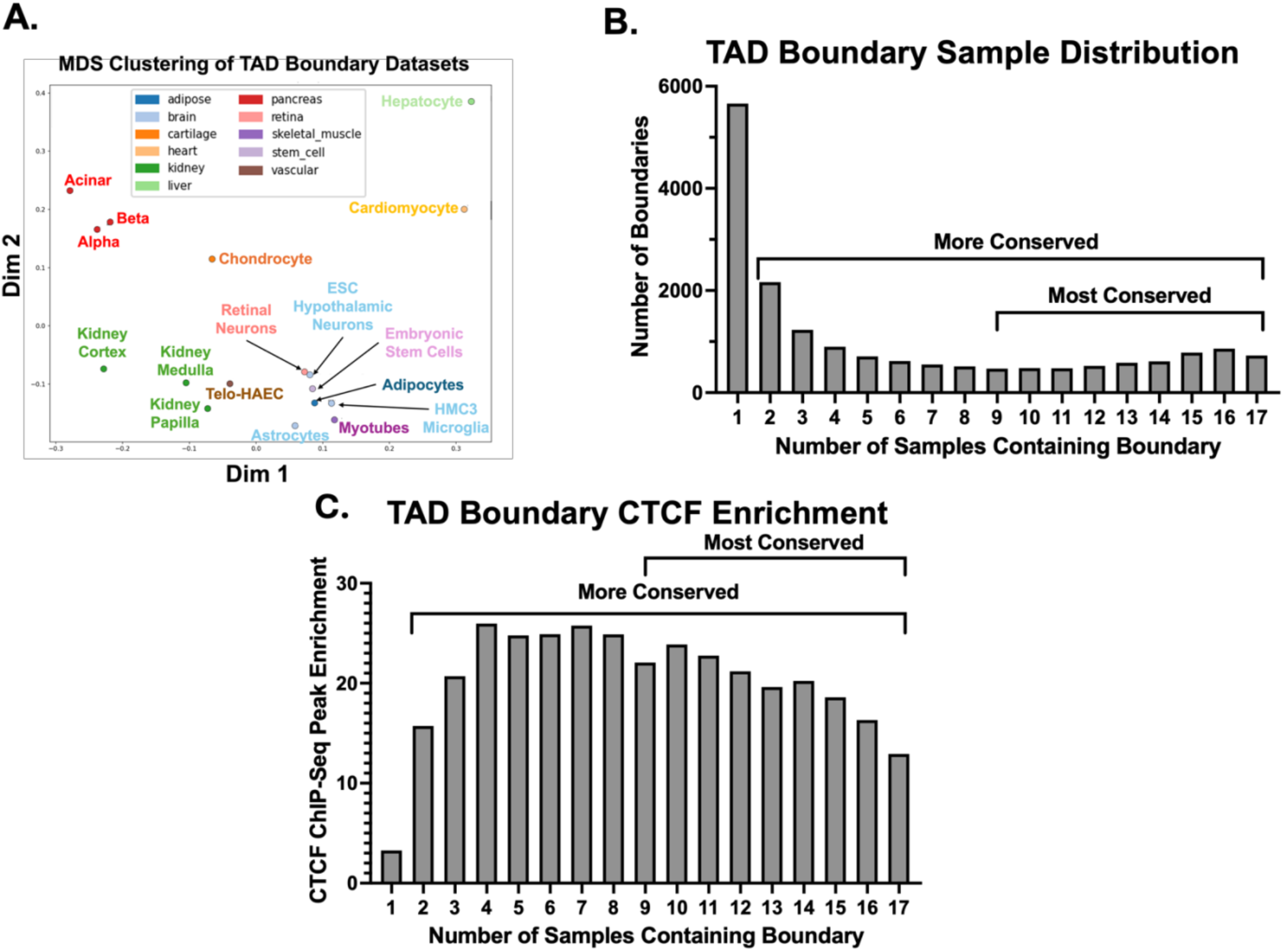
Topologically associated domain (TAD) boundary dataset analysis. **A**. MDS clustering of TAD boundary locations indicates tissue specific clustering. **B.** Number of TAD boundaries that are shared across samples. Many boundaries are unique to one sample. Distribution shows increasing boundary numbers as the sharing requirement increases, indicating a population of boundaries that are conserved across tissues (labelled “most conserved”). **C.** Analysis of CTCF ChIP-Seq signal overlapping boundaries. Boundaries present in a single sample show modest CTCF signal enrichment, which supports their interpretation as noise. Conserved boundaries show higher CTCF enrichment scores, supporting their role as conserved TAD boundaries. TAD, topologically associated domain. MDS, multidimensional scaling plot.

**Supplemental Table 1.**
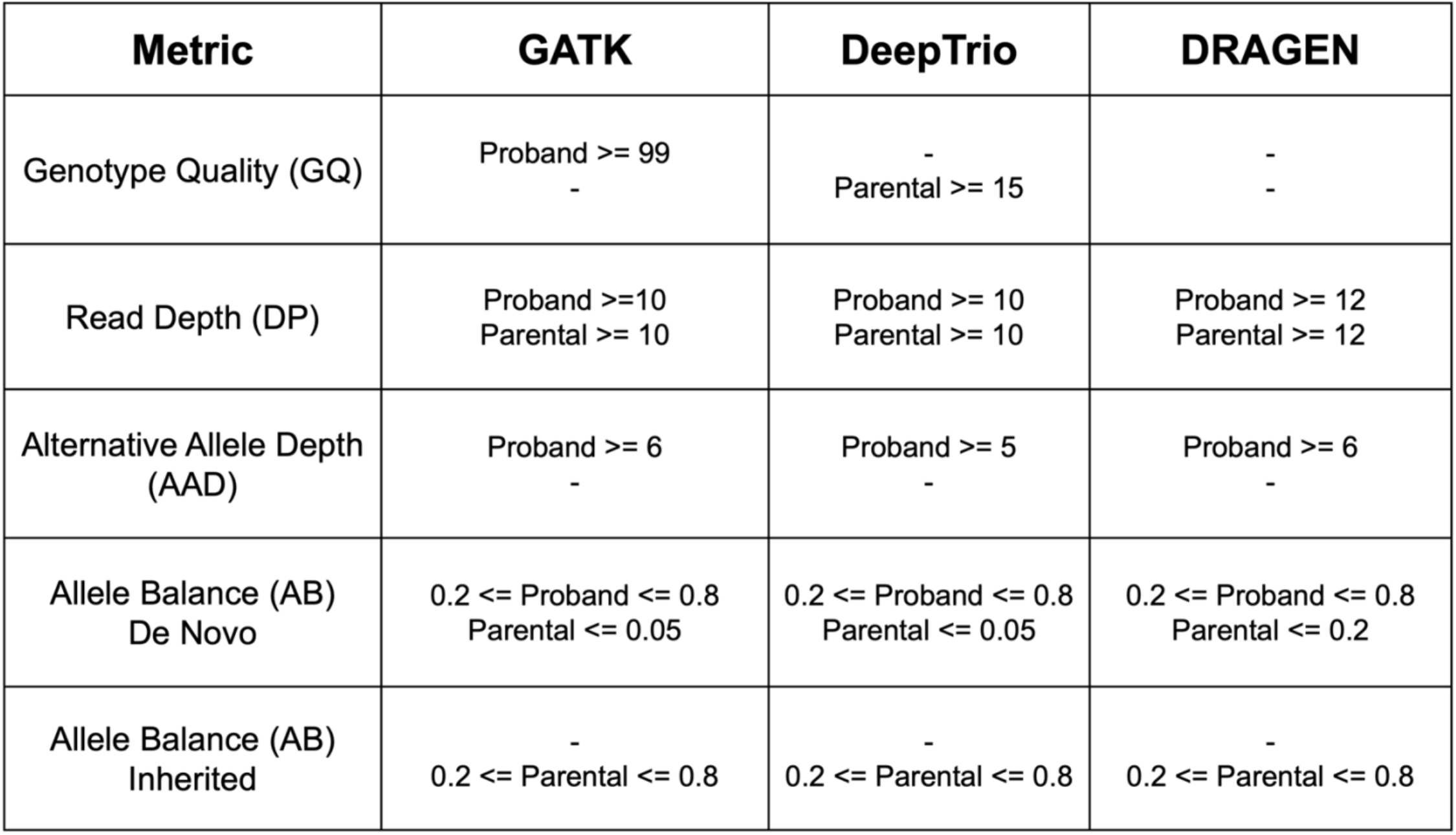
Variant quality control filtering metrics determined by GIAB benchmarking. QC cutoffs were determined by comparing the distribution of true positive and false positive de novo variants from GIAB. Cutoffs were set to maximize recall at the expense of decreased precision.

**Supplemental Table 2. All de novo and rare inherited variants called by the GAVURD variant calling pipeline on 10 unsolved rare disease probands.**

**Supplemental Table 3. Arima Hi-C datasets used for TAD boundary analysis.**

**Supplemental Table 4. Human Phenotype Ontology terms for 10 unresolved rare disease probands.**

**Supplemental Table 5. Top 10 candidate causative non-coding variants found in 10 unresolved rare disease probands.**

